# Paradoxical excitation of lateral habenula neurons by propofol involves enhanced presynaptic release of glutamate

**DOI:** 10.1101/2021.11.08.467738

**Authors:** Ryan David Shepard, Kunwei Wu, Wei Lu

## Abstract

Sleep is a fundamental physiological process conserved across most species. As such, deficits in sleep can result in a myriad of psychological and physical health issues. However, the mechanisms underlying the induction of sleep are relatively unknown. Interestingly, general anesthetics cause unconsciousness by positively modulating GABA-A receptors (GABA_A_Rs). Based on this observation, it is hypothesized that GABA_A_Rs play a critical role in modulating circuits involved in sleep to promote unconsciousness. Recently, the lateral habenula (LHb) has been demonstrated to play a role in sleep physiology and sedation. Specifically, propofol has been shown to excite LHb neurons to promote sedation. However, the mechanism by which this occurs is unknown. Here, we utilize whole-cell voltage and current clamp recordings from LHb neurons obtained from 8-10 week old male mice to determine the physiological mechanisms for this phenomenon. We show that bath application of 1.5μM propofol is sufficient to increase LHb neuronal excitability involving synaptic transmission, but not through modulation of intrinsic properties. Additionally, although there is increased LHb neuronal excitability, GABA_A_Rs localized postsynaptically on LHb neurons are still responsive to propofol, as indicated by an increase in the decay time. Lastly, we find that propofol increases the synaptic drive onto LHb neurons involving enhanced presynaptic release of both glutamate and GABA. However, the greatest contributor to the potentiated synaptic drive is the increased release of glutamate which shifts the balance of synaptic transmission towards greater excitation. Taken together, this study is the first to identify the physiological basis for why LHb neurons are excited by propofol, rather than inhibited, and as a result promote sedation.

## Introduction

Sleep is an indispensable physiological process that is evolutionarily conserved and is vital for maintaining overall health. Not surprisingly, sleep deficits predispose individuals to major health issues including obesity, diabetes, cardiovascular issues and psychiatric conditions [1]. Indeed, a lack of sleep has been shown to contribute to impaired learning and memory [2,3], as well as a higher comorbidity for anxiety and mood disorders [4,5]. However, determining the importance of sleep and elucidating the mechanisms underlying sleep induction, as well as maintenance, still remains relatively unknown which unfortunately has translated to little efficacious therapies devoid of unwanted side effects.

Various molecular targets ranging from neurotransmitters to neuropeptides have been postulated to be involved with dampening arousal circuits and enhancing the overall activity of sleep-promoting regions which has generated a variety of hypothetical models [6,7]. Of interest, is the role of inhibition through GABAergic circuits that exert their fast inhibitory effects through GABA_A_Rs, pentameric ligand-gated ion channels differentially assembled from 19 subunits that endow discrete biophysical and pharmacological properties [8,9]. GABA_A_R regulation consists of dynamic processes involving a myriad of intracellular mechanisms, scaffolds, and more recently novel transmembrane proteins [10–13]. Given their diversity and ubiquity across the brain, GABA_A_Rs have been long sought as targets for various drug classes such as for treatment of epilepsy, anxiety, depression, and sleep disorder [9]. Indeed, general anesthetics (GAs), benzodiazepines, nonbenzodiazepines (also known as “Z-drugs”), and neurosteroids all target GABA_A_Rs and can induce sedation despite their different binding sites on GABA_A_Rs, suggesting that GABA_A_R activation is a universal mechanism involved with promoting sleep states.

Recently, studies have shown that GAs, such as propofol [14,15], and isoflurane [16], have a pronounced effect on the lateral habenula (LHb), an epithalamic structure which has been implicated in various psychiatric disorders such as substance abuse disorder, sleep disorder, and most notably depression [17]. The LHb serves as a critical interface between forebrain, as well as limbic regions, and monoaminergic structures by receiving dense inputs via the stria medullaris from areas such as medial prefrontal cortex and globus pallidus (rodent homolog is the entopeduncular nucleus). Because glutamatergic LHb neurons potently regulate monoamines (such as ventral tegmental area dopamine [DA] neurons [18–20]) by glutamatergic efferents via the fasciculus retroflexus, it is not surprising that the LHb is poised to also regulate sedation given the role of DA in arousal. Indeed numerous studies have suggested that the LHb potentially regulates sleep [21] through evidence demonstrating that the structure has intrinsic circadian activity [22,23], is responsive to zeitgebers like light [22–24], and that the structure receives inputs from other sleep-associated regions such as the lateral hypothalamus (LH), basal forebrain, lateral preoptic area, and suprachiasmatic nucleus [25]. Mechanistically, it is still not well understood how modulation of LHb activity regulates sleep, but pharmacologically it appears GABA_A_R activation is a contributing mechanism. However, to date, there are a paucity of studies examining the physiological significance and impact of pharmacological manipulation on GABA_A_Rs within the LHb. Given that propofol increases LHb activity to induce sedation [14,15], here we utilized *in vitro* electrophysiological recordings from mice LHb neurons to determine the physiological mechanism for this phenomenon.

## Results

Previous studies have demonstrated propofol exposure elevates c-fos immunoreactivity within the LHb [14,15,26]. Additionally, it was previously reported that propofol (1.5μM) does not affect resting membrane potential (RMP) of LHb neurons [14]. Using whole-cell current clamp recordings and a 1.5μM concentration of propofol, we confirmed this previous finding (**Figure 1A**) and also expanded upon this observation demonstrating that regardless of the firing pattern (silent, tonic or burst; [17]), there are no differences in RMP of cells recorded in the absence or presence of propofol (**Figure 1B**). Next, we challenged LHb neurons with stepwise injections of increasingly depolarizing current to quantify LHb neuronal excitability in the absence or presence of propofol. Interestingly, propofol-induced increases in LHb neuronal excitability only occurred when synaptic transmission was intact (**Figure 1C-E**). In the absence of synaptic transmission, propofol did not affect RMPs or intrinsic neuronal excitability (**Figure 2**). Additionally, at our concentration of propofol, AP half-width was decreased when synaptic transmission was intact (**Supplementary Table 1**) which was not observed in the absence of fast synaptic transmission (**Supplementary Table 2**). Taken together, it would appear that propofol utilizes synaptic transmission to enhance LHb neuronal excitability.

**Figure 1.**
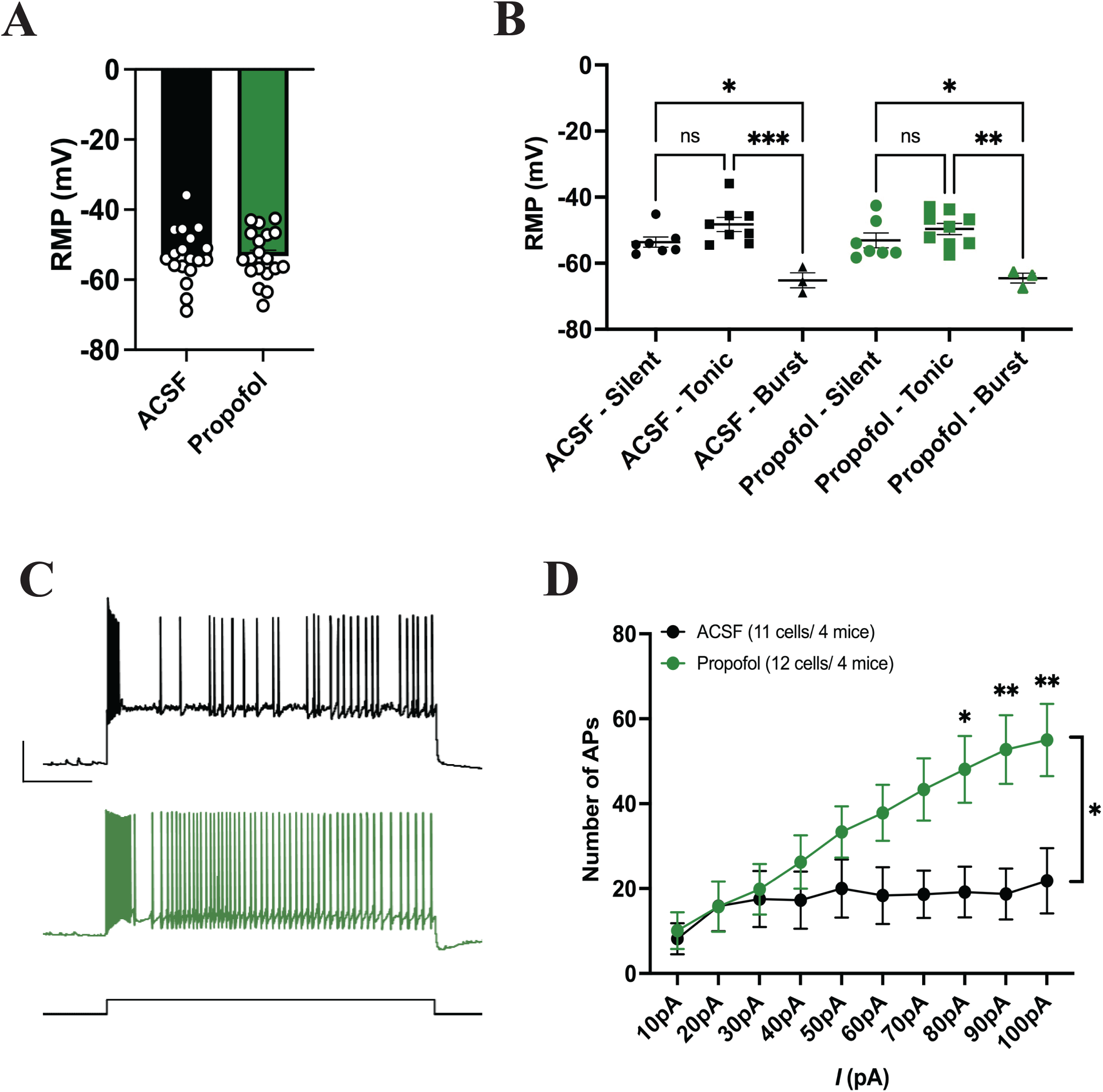
Propofol increases LHb neuronal excitability. **A**. Propofol does not affect the overall RMP of LHb neurons (ACSF, n=18 cells/7 mice; Propofol, n=19 cells/7 mice; 2-tailed independent t-test, t(35)=0.04267, P=0.9662). **B**. Propofol does not affect the RMP of LHb neurons in a cell-type specific manner (ACSF, n=18 cells/7 mice; Propofol, n=19 cells/7 mice; one-way ANOVA, F(5,31)=8.578, P<0.0001. **C**. Representative excitability traces of neurons recorded in ACSF or ACSF + Propofol at a current step of 20pA. Calibration bars: 20mV/1s. **D**. Propofol increases the number of action potentials produced by LHb neurons with increasing steps of depolarizing current (ACSF, n=11 cells/4 mice; Propofol, n=12 cells/4 mice; two-way RM ANOVA, Current: F(9,189) = 12,57, P<0.001; Drug: F(1,21)=4.565, P=0.0446; Current x Drug: F(9,189)=6.193, P<0.001.

**Figure 2.**
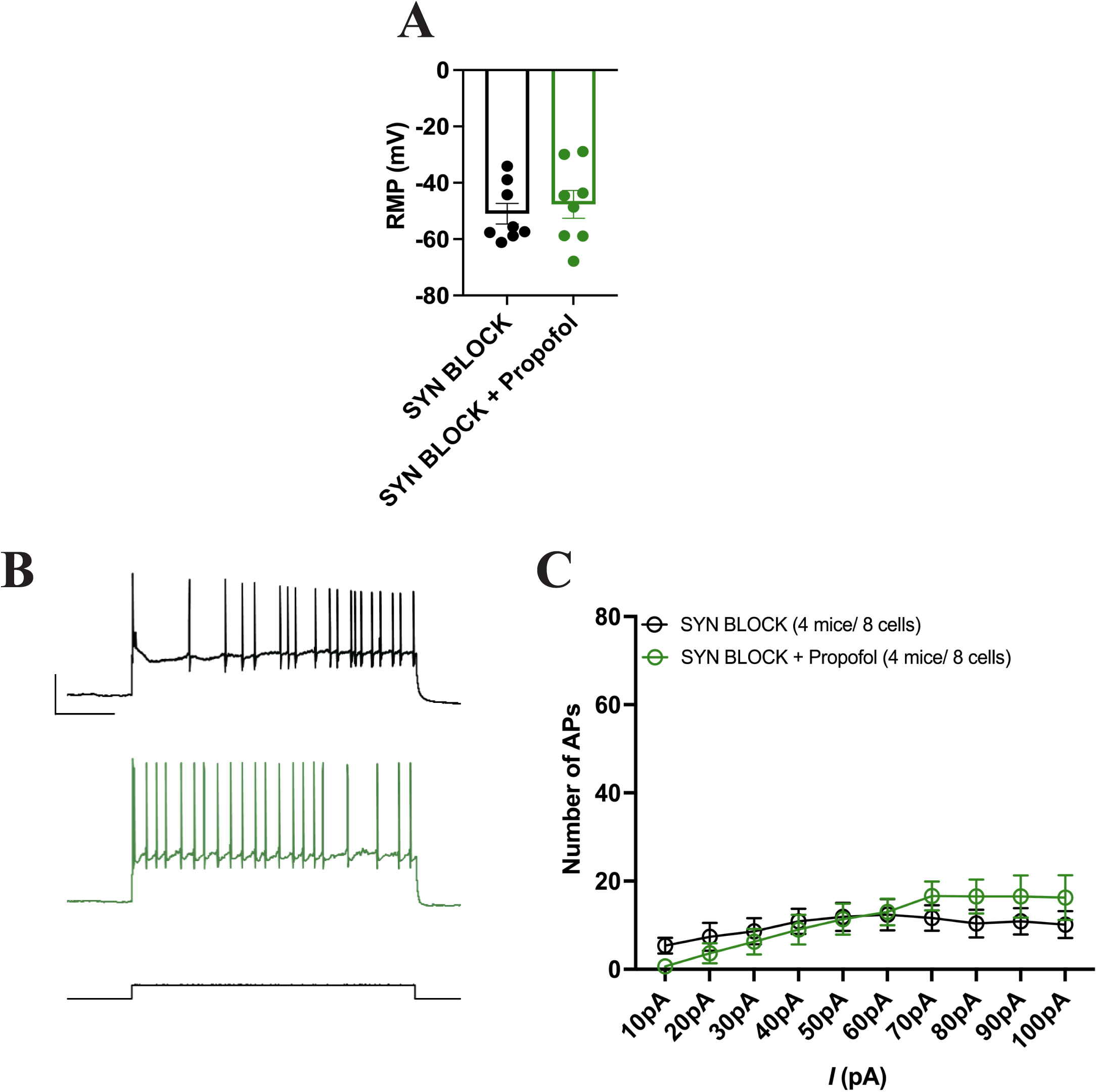
Propofol does not affect LHb intrinsic neuronal excitability when fast synaptic transmission is blocked. **A**. Propofol does not affect the overall RMP of LHb neurons (SYN BLOCK, n=8 cells/4 mice; SYN BLOCK + Propofol, n=8 cells/4 mice; 2-tailed independent t-test, t(14)=0.5460, P=0.5937). **B**. Representative excitability traces of neurons in the presence of antagonists for fast synaptic transmission (APV, DNQX, PTX) or APV + DNQX + PTX + Propofol at a current step of 20pA. Calibration bars: 20mV/1s. **C**. Propofol does not affect the number of action potentials by LHb neurons intrinsically with increasing steps of depolarizing current (SYN BLOCK, n=8 cells/4 mice; SYN BLOCK + Propofol, n=8 cells/4 mice; two-way RM ANOVA, Current F(2.190, 30.66)=5.141, P=0.0101; Drug: F(1,14)=0.1015, P=0.7574; Current x Drug: F(9,126)=1.582, P=0.1274).

Based on our findings that propofol-induced increases in LHb neuronal excitability require synaptic transmission, we examined whether spontaneous excitatory and inhibitory were affected. Compared to baseline, neither the frequency or amplitude of spontaneous excitatory postsynaptic currents (sEPSCs) were affected by propofol (**Figure 3A-C**). Interestingly, propofol increased the frequency of spontaneous inhibitory postsynaptic currents (sIPSCs) without affecting amplitude (**Figure 3D-F**). However, the decay of sIPSCs was significantly prolonged indicating that GABA_A_Rs sitting postsynaptically on LHb neurons respond to propofol (**Figure 3G**). It is important to note that under this experimental condition, isolating AMPAR-mediated sEPSCs requires applying an antagonist (picrotoxin) that blocks GABA_A_Rs. To overcome this, we recorded postsynaptic currents using a low chloride internal which allowed for simultaneous acquisition of both AMPAR- and GABA_A_R-mediated currents from LHb neurons by voltage-clamping neurons at −40mV as done previously [27,28]. By allowing propofol to positively modulate GABA_A_Rs throughout the slice, we observed enhanced synaptic drive which was driven by both increased sEPSC and sIPSC frequency; however, the sEPSC frequency was potentiated to a greater extent (**Figure 4**). Collectively, GABA_A_Rs within the LHb are responsive to propofol, but this is not sufficient to overcome propofol-induced increases in LHb neuronal excitability due to increased glutamate release presynaptically.

**Figure 3.**
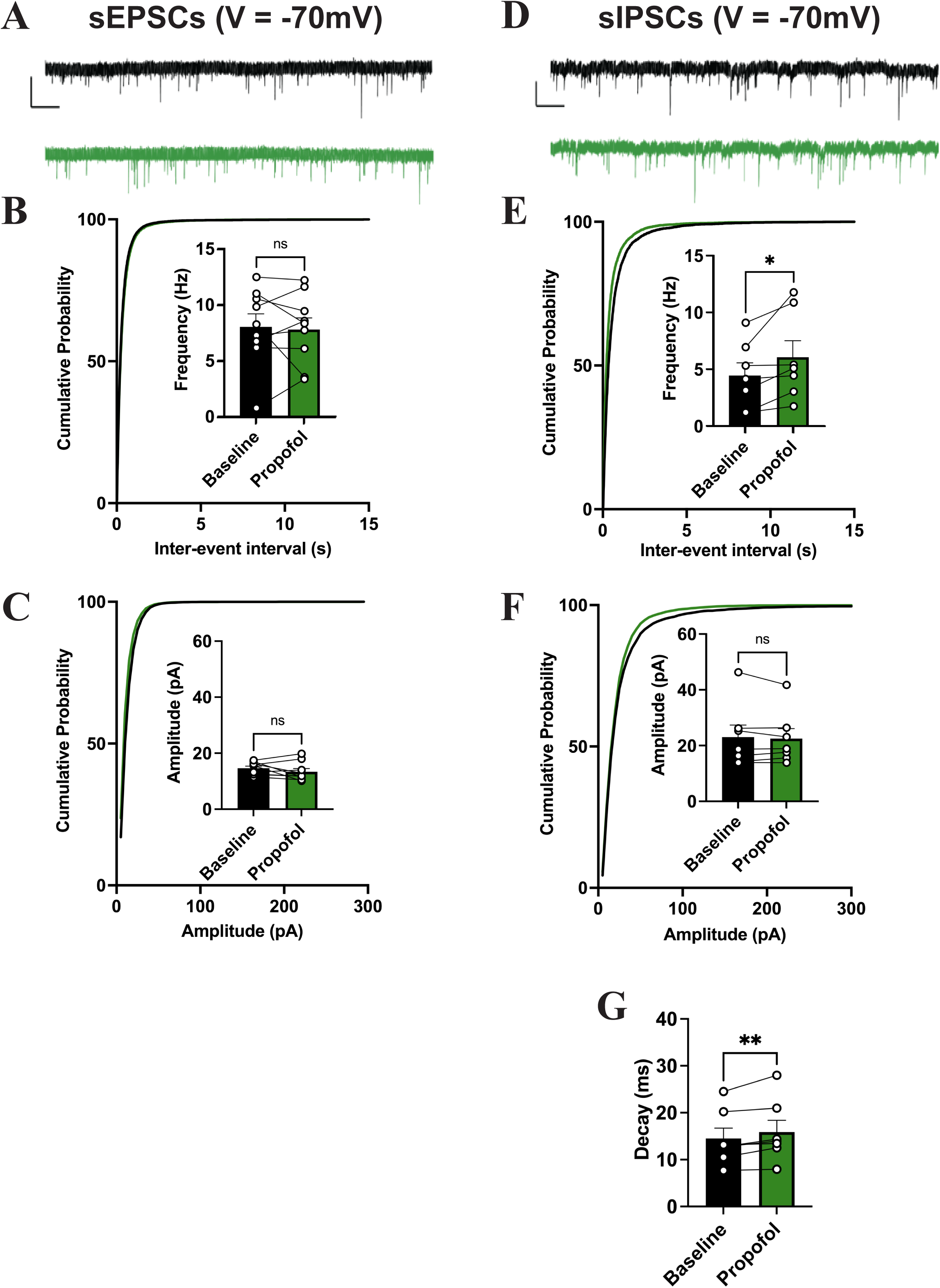
Effects of propofol on spontaneous excitatory and inhibitory postsynaptic currents. **A**. Representative trace of AMPAR-mediated sEPSCs obtained from LHb neurons at a holding potential of −70mV. Calibration bars: 20pA/200ms. **B**. and **C**. Propofol does not affect the frequency (Baseline and Propofol n=9 cells/7 mice; one-tailed paired t-test, t(8)=0.3144, P=0.3806) or amplitude of sEPSCs (Baseline and Propofol n=9 cells/7 mice; one-tailed paired t-test, t(8)=1.268, P=0.1202). **D**. Representative trace of GABA_A_R-mediated sIPSCs obtained from LHb neurons at a holding potential of −70mV. Calibration bars: 20pA/200ms. **E**. and **F**. Propofol increases the frequency of sIPSCs (Baseline and Propofol n=7 cells/4 mice; one-tailed paired t-test, t(6)=2.637, P=0.0193), but does not alter the amplitude (Baseline and Propofol n=7 cells/4 mice; one-tail paired t-test, t(6)=0.6677, P=0.2646). **G**. Propofol enhances the sIPSC decay time (Baseline and Propofol n=7 cells/4 mice; one-tailed paired t-test, t(6)=3.211, P=0.0092). P*< 0.05, P**< 0.01.

**Figure 4.**
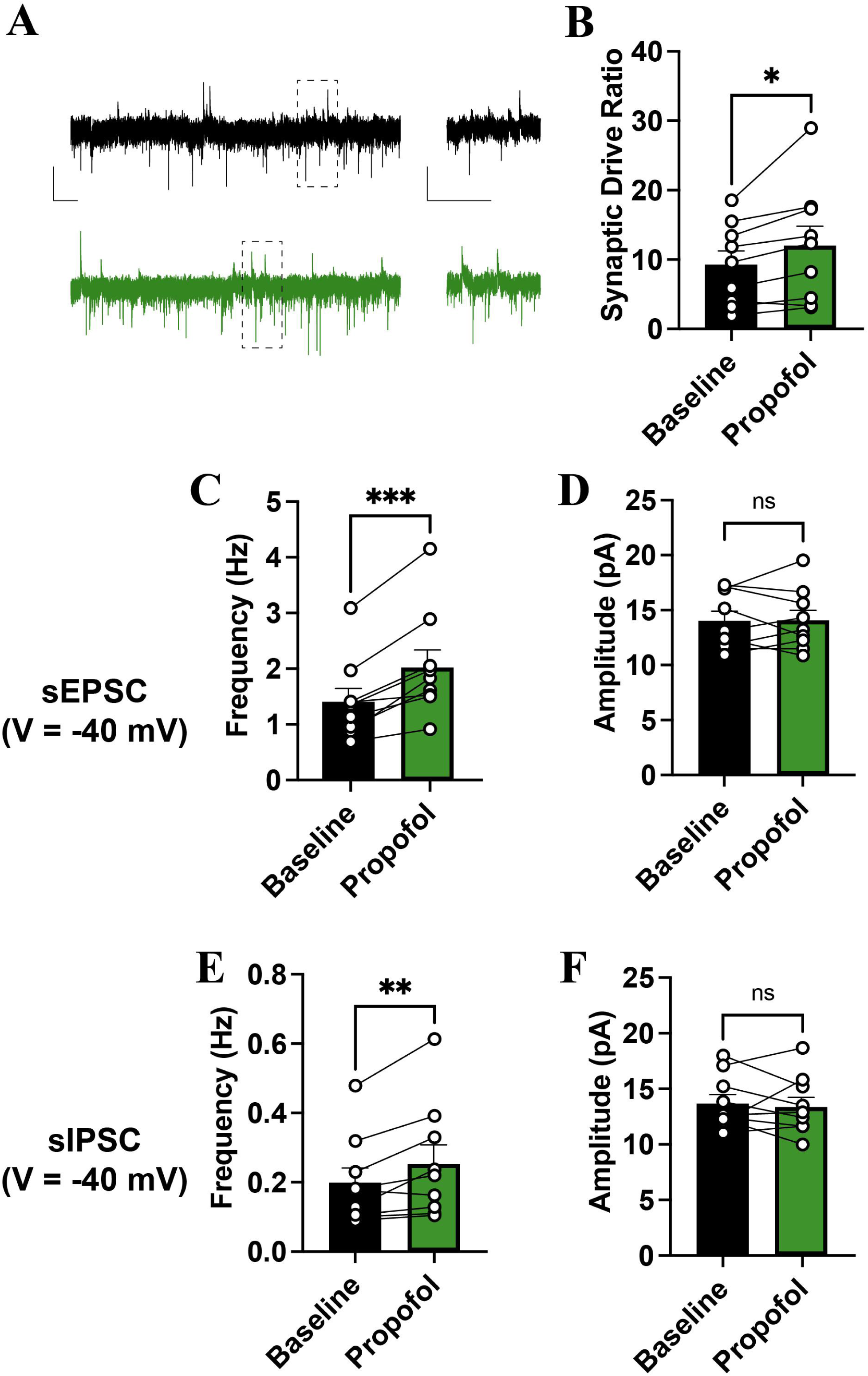
Propofol alters the balance of presynaptic release of glutamate/GABA by enhancing glutamatergic transmission. **A**. Representative sPSC traces at V=-40mV using low chloride internal (downward deflections are AMPAR-mediated EPSCs and upward deflections are GABA_A_R-mediated IPSCs) from LHb neurons. Calibration bars: 20pA/2s. **B**. Propofol increases the synaptic drive ratio ([mean EPSC frequency x mean EPSC amplitude]/[mean IPSC frequency x mean IPSC amplitude]) on LHb neurons (baseline, n=9 cells/5 mice; propofol, n=9 cells/5 mice; one-tailed paired t-test; t(8)=2.638, p = 0.0149). **C**. and **D**. Propofol enhances sEPSC frequency (baseline, n=9 cells/5 mice; propofol, n=9 cells/5 mice; one-tailed paired t-test; t(8)=5.662, p = 0.0002) while not affecting sEPSC amplitude (baseline, n=9 cells/5 mice; propofol, n = 9 cells/5 mice; one-tailed paired t-test; t(8) = 0.07309, p = 0.4718). **E**. and **F**. Propofol enhances sIPSC frequency (baseline, n = 9 cells/5 mice; propofol, n=9 cells/5 mice; one-tailed paired t-test; t(8)=0.3148, p=0.0068) while not affecting amplitude (baseline, n=9 cells/5 mice; propofol, n=9 cells/5 mice; one-tailed paired t-test; t(8)=0.4532, P=0.3312). P*< 0.05, P** < 0.01, P*** < 0.001.

## Discussion

To our knowledge, this is the first study to mechanistically determine why propofol increases LHb neuronal excitability which seems critical for its sedative properties [14]. In line with previous observations [14,15] and based on our findings of propofol-induced enhancement of sEPSCs when GABA_A_Rs are not blocked by picrotoxin, disinhibition of upstream excitatory inputs could be a plausible mechanism underlying the increased LHb neuronal excitability. Additionally, this also could potentially contribute to our observed decrease in the AP half-width given that decreased half-width is a property associated with a greater frequency of AP production [29]. However, the exact upstream circuits mediating this effect are most likely complex and could potentially involve multiple upstream structures. One study examining circuit-based mechanisms for isoflurane sedation demonstrated that a potential input are glutamatergic neurons originating from the LH [30]. Although there has been much quandary over potential local GABAergic neurons within the LHb [31–34], it is interesting to speculate whether disinhibition of LHb neurons directly involves this subpopulation, although this still requires intensive and further validation for a physiological role for LHb GABAergic neurons.

It is also worth noting that a majority of studies have focused on the mechanistic role of AMPARs and NMDARs in regulating LHb physiology, however there are a paucity of studies regarding how GABA_A_Rs are involved. Unlike other structures where exposure to propofol decreases neuronal excitability [35,36], this does not seem to be the case here. Interestingly, direct intra-LHb injections of GABAergic agonists and antagonists have been shown to decrease and increase LHb activity [37], respectively, indicating that GABA_A_Rs do indeed serve an inhibitory role within the LHb. This is also supported by dense KCC2 expression [38] allowing for low intracellular chloride concentrations within LHb neurons. Given that we observed an increase in sIPSC decay time, it is unlikely that the pharmacological properties underlying GABA_A_R function are different from what has been observed canonically. Although we found that propofol did not affect the RMP of LHb neurons, this could be due to a variety of factors such as low ambient GABA. With respect to our recordings of synaptic transmission, the main difference in why we detected increased sEPSC frequency when recording simultaneous sEPSCs and sIPSCs compared to sEPSCs in isolation is due to experimental conditions. By recording dual currents without blocking GABA_A_Rs, we allowed for potential upstream structures to be modulated (e.g., disinhibition) and were then able to observe enhanced sEPSC frequency. In addition to this being the first study to demonstrate how propofol affects the cellular physiology of LHb neurons, the only other GABA_A_R-targeting drug which has been mechanistically evaluated in the LHb is ethanol, which interestingly also promotes increased LHb excitability involving presynaptic mechanisms [27].

One limitation of our study is that we only utilized 8-10 week old male mice to remain consistent with a previous study that showed propofol-induced activation did not alter the RMP of LHb neurons [14] at 1.5μM. Due to the fact that hormones can impact sleep, it is important to examine whether the effect of propofol that we and others have observed on LHb neurons is conserved in females. With respect to the chosen propofol concentration for our experiments, it remains to be seen if all propofol concentrations capable of inducing sedation involve increased glutamate release onto LHb neurons. Additionally, propofol has also been shown to affect neuronal excitability through SK [39] and HCN channels [40], both of which can regulate LHb neurophysiology and AP properties [41,42]. In order to more comprehensively evaluate the effect of propofol and other GAs on LHb physiology, further examination of the physiological response of LHb neurons to differential concentrations of GAs is warranted. Lastly, our recordings took place during “lights on” and it has been shown at least for mEPSCs that presynaptic release probability is affected temporally based on the time of day [43] and therefore it would be interesting to see 1) if IPSCs are also affected temporarily and 2) whether the robustness of LHb response to propofol is circadian dependent.

## Materials and Methods

### Animals

Animal handling was conducted in accordance with all rules, regulations, and protocols approved by the NIH/NINDS Institutional Animal Care and Use Committee (IACUC). 8-10 week male C567BL/6N purchased from Charles River were used for slice electrophysiology experiments. All mice were housed in a conventional vivarium on a 12-hour light/dark cycle and were allowed ad libitum access to food and water. All efforts were made to minimize the number of mice used and to reduce any animal suffering. Mice noted as unhealthy by the veterinarian or veterinarian staff were excluded from the study.

### Slice Preparation

Mice were anesthetized by isoflurane and were confirmed unresponsive by pinching the back paw to determine whether there was any withdrawal reflex. Mice brains were quickly dissected in ice-cold artificial cerebrospinal fluid (ACSF) containing (in mM): 126 NaCl, 21.4 NaHCO3, 2.5 KCl, 1.2 NaH2PO4, 2.4 CaCl2, 1.00 MgSO4, 11.1 glucose, 0.4 ascorbic acid, saturated with 95% O2–5% CO2. Sagittal and coronal brain slices containing the LHb were cut at 300um and incubated in ACSF at 34C for at least 90 minutes. Slices were transferred to recording chambers and perfused with ACSF lacking ascorbic acid at 28C.

### Electrophysiology

Whole cell recordings were performed on slices containing LHb using Multiclamp 700B under infrared-differential interference contrast microscopy. Data acquisition was carried out using both DigiData 1550b1 and (insert acquisition board here). Recordings were acquired at 10kHz and filtered at 3Hz. Series (5-25 MΩ) was monitored throughout the recording and recordings were excluded from the study if series value deviated by > 25% during the recording. Whole-cell recordings were used in order to determine the predominant firing pattern of LHb neurons (tonic, burst, silent) and resting membrane potential (RMP) in current clamp without injecting current (i.e., I=0). Neuronal excitability whole-cell recordings on LHb neurons were conducted as previously described [41,42,44]. LHb current clamp recordings were conducted using a potassium gluconate based internal containing: 130 mM K-gluconate, 15 mM KCl, 4 mM ATP-Na+, 0.3 mM GTP-Na+, 1 mM EGTA, and 5 mM HEPES (pH adjusted to 7.28 with KOH, osmolarity adjusted to 280–290 mOsm). LHb neurons were manually injected with sufficient current to maintain a holding membrane potential of ∼ −68mV. Increasing steps of depolarizing current of +10pA current (from 10pA to 100pA) for a duration of 5 seconds. Current injections were separated by a 20 second interval. The number of APs produced during depolarization were quantified. Any APs that occurred outside of the 5 second current injection period were not included. Membrane properties including fast afterhyperpolarization (fAHP), medium afterhyperpolarization (mAHP), action potential threshold (AP threshold), action potential amplitude (AP amplitude), and input resistance (Rin) were measured as described in detail in [41]. Spontaneous excitatory postsynaptic currents (sEPSCs) and spontaneous inhibitory postsynaptic currents (sIPSCs) were conducted using whole-cell voltage clamp recordings by clamping LHb neurons at −70mV. AMPAR-mediated sEPSCs were recorded using a cesium gluconate internal containing the following: 117 mM Cs-gluconate, 2.8 mM NaCl, 5 mM MgCl2, 2 mM ATP-Na+, 0.3 mM GTP-Na+, 0.6 mM EGTA, and 20 mM HEPES (pH adjusted to 7.28 with CsOH, osmolarity adjusted to 275–280 mOsm). The recording solution (ACSF) contained picrotoxin (100µM) and D-APV (50µM) to block GABA_A_Rs/GlyRs and NMDARs, respectively. GABA_A_R-mediated sIPSCs were recorded using a cesium chloride (high chloride internal) containing in mM: 130 CsCl, 8.5 NaCl, 5 HEPES, 4 MgCl2, 4 Na-ATP, 0.3 Na-GTP and 1 QX-314 pH 7.2. ACSF contained DNQX, APV, and strychnine to block AMPARs, NMDARs, and GlyRs, respectively. Dual PSCs were assayed as previously described [27,28] using the following recording solution in mM: 126 NaCl, 2.5 KCl, 1.25 NaH2PO4, 1 MgCl2, 2 CaCl2, 25 NaHCO3, 0.3 l-ascorbate, and 11 glucose. A cesium methylsulfanate (low chloride internal) was used containing: 140 Cs-methanesulfonate, 5 KCl, 2 MgCl2, 10 HEPES, 2 MgATP, 0.2 NaGTP, pH 7.2. Dual PSCs were recorded at −40mV which was sufficient to collect AMPAR-mediated sEPSCs as downward current deflections and GABA_A_R-mediated sIPSCs as upward current deflections. Electrophysiological recordings were analyzed off-line utilizing ClampFit 11, MiniAnalysis, and IgorPro 6.22A. ClampFit 11 was used to analyze current clamp recordings. MiniAnalysis and IgorPro 6.22A were used to analyze voltage clamp recordings. sEPSCs and sIPSCs were detected using pre-specified parameters with an amplitude cutoff of 5pA.

### Statistical Analysis

All values are presented as the mean +/-the standard error of the mean (S.E.M.). All data sets were tested for normality and tested using the corresponding parametric and non-parametric tests. Statistical comparisons used included independent t-test, dependent t-test, Mann-Whitney U, one-way ANOVA and two-way repeated measure ANOVA (RM-ANOVA). If ANOVAs were significant, Bonferroni post hoc analysis was utilized. All comparisons had a threshold set for significance at p<0.05. Statistical analysis was conducted using GraphPad Prism 9.

## Funding

This work was supported by the NIH/NINDS Intramural Research Program (to W.L.) and a Postdoctoral Fellowship from the NIH Center on Compulsive Behaviors (to R.D.S.).

## Author Contributions

R.D.S. designed the project and W.L. supervised the project. R.D.S. and K.W. performed electrophysiology experiments. R.D.S., K.W., and W.L. wrote the manuscript, and all authors read and commented on the manuscript.

## Disclosures

The authors declare that no competing interests exist.

## Figure Legends

**Supplementary Figure 1.**
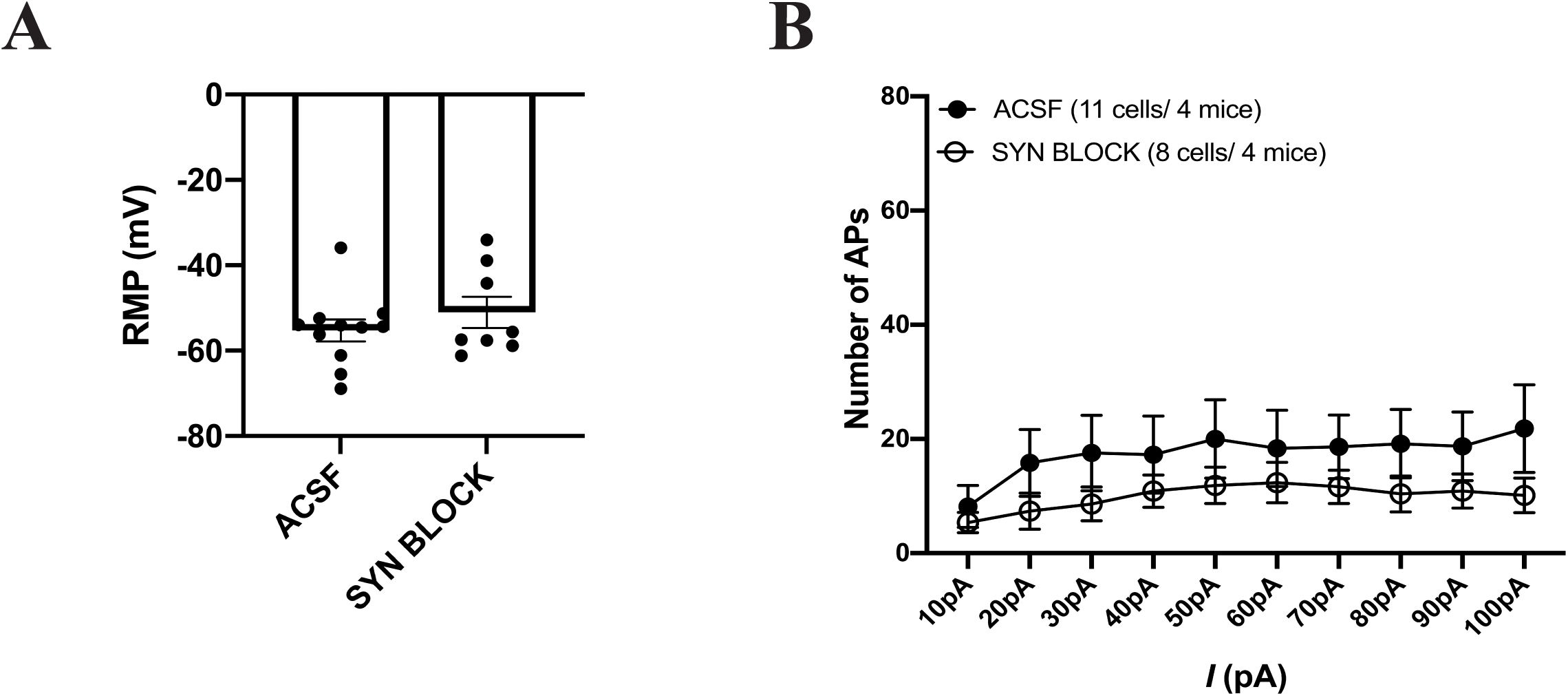
Blockade of fast synaptic transmission does not significantly alter the RMP or neuronal excitability of LHb neurons. **A**. RMP of LHb neurons are not significantly affected in the absence of fast synaptic transmission (ACSF, n=11 cells/4 mice; SYN BLOCK, n=8 cells/4 mice; two-tailed unpaired t-test; t(18)=0.9915, P=0.3353). **B**. Block of fast synaptic transmission (AMPARs, NMDARs, and GABA_A_Rs) does not significantly affect LHb neuronal excitability (ACSF, n=11 cells/4 mice; SYN BLOCK, n=8 cells/4 mice; two-way RM ANOVA, Current: F(9,153) = 2.016, P<0.0409; Treatment: F(1,17)=1.232, P=0.2825; Current x Treatment: F(9,153)=0.3473, P=0.9574).

**Supplementary Table 1**. <u>Summary of action potential properties of neurons exposed to ACSF or Propofol.</u>

All values are means ± SEM. **\***P<0.05, independent t-test. Abbreviations: AP, action potential; fAHP, fast afterhyperpolarization; mAHP, medium afterhyperpolarization; Rin, input resistance.

**Supplementary Table 2**. <u>Summary of action potential properties of neurons when fast synaptic transmission was blocked or when fast synaptic transmission was blocked and Propofol.</u>

Values were extracted from neurons reported in **Figure 1B**. All values are means ± SEM. N.S., independent t-test. Abbreviations: AP, action potential; fAHP, fast afterhyperpolarization; mAHP, medium afterhyperpolarization; Rin, input resistance.

**Supplementary Table 3**. <u>Summary of action potential properties of neurons exposed to ACSF or when fast synaptic transmission was blocked.</u>

Values were extracted from neurons reported in **Supplementary Figure 1**. All values are means ± SEM. **\*\***P<0.01, independent t-test. Abbreviations: AP, action potential; fAHP, fast afterhyperpolarization; mAHP, medium afterhyperpolarization; Rin, input resistance; SYN BLOCK, synaptic blockade.

---

**Supplementary Table 1.** Summary of AP properties and membrane properties with synaptic transmission intact of LHb neurons either exposed to bath-applied drug-free ACSF or ACSF containing propofol (1.5µM). Values were extracted from neurons reported in **Figure 1A**. All values are means ± SEM. **\***P < 0.05, independent t-test. Abbreviations: AP, action potential; fAHP, fast afterhyperpolarization; mAHP, medium afterhyperpolarization; Rin, input resistance.

| Drug Treatment<br>( # of cells/# of animals) | ACSF (11/4) | Propofol (12/4) |
| --- | --- | --- |
| AP threshold (mV) | -38.58 ± 1.868 | -40.06 ± 2.148 |
| AP peak (mV) | 97.42 ± 1.684 | 96.29 ± 3.000 |
| AP half width (ms) | 1.693 ± 0.1589 | 1.291 ± 0.09547 * |
| fAHP (mV) | -12.30 ± 1.807 | -10.78 ± 1.567 |
| mAHP (mV) | -35.28 ± 1.994 | -32.83 ± 2.463 |
| Rin (mΩ) | 386.3 ± 62.24 | 368.7 ± 43.61 |

**Supplementary Table 2.** Summary of AP properties and membrane properties with synaptic transmission blocked of LHb neurons either exposed to bath-applied ACSF containing APV (50 µM), DNQX (10 µM), PTX (100µM) or ACSF containing APV (50 µM), DNQX (10 µM), PTX (100µM), and propofol (1.5µM). Values were extracted from neurons reported in **Figure 1B**. All values are means ± SEM. N.S., independent t-test. Abbreviations: AP, action potential; fAHP, fast afterhyperpolarization; mAHP, medium afterhyperpolarization; Rin, input resistance.

| Drug Treatment<br>( # of cells/# of animals) | SYN BLOCK (11/4) | SYN BLOCK + Propofol (12/4) |
| --- | --- | --- |
| AP threshold (mV) | -34.83 ± 1.993 | -35.88 ± 1.446 |
| AP peak (mV) | 106.5 ± 2.482 | 103.5 ± 3.933 |
| AP half width (ms) | 2.019 ± 0.1475 | 1.562 ± 0.1762 |
| fAHP (mV) | -13.16 ± 1.994 | -10.45 ± 0.6937 |
| mAHP (mV) | -38.25 ± 2.320 | -38.43 ± 1.660 |
| Rin (mΩ) | 495.6 ± 68.72 | 401.6 ± 44.33 |

**Supplementary Table 3.** Summary of AP properties and membrane properties with synaptic transmission blocked of LHb neurons either exposed to bath-applied ACSF or ACSF containing APV (50 µM), DNQX (10 µM), PTX (100µM). Values were extracted from neurons reported in **Supplementary Figure 1**. All values are means ± SEM. **\*\***P < 0.01, independent t-test. Abbreviations: AP, action potential; fAHP, fast afterhyperpolarization; mAHP, medium afterhyperpolarization; Rin, input resistance; SYN BLOCK, synaptic blockade.

| **Drug Treatment**<br>**( # of cells/# of animals)** | **ACSF (11/4)** | **SYN BLOCK (8/4)** |
| --- | --- | --- |
| AP threshold (mV) | -38.58 ± 1.868 | -34.83 ± 1.993 |
| AP peak (mV) | 97.42 ± 1.684 | 106.5 ± 2.482** |
| AP half width (ms) | 1.693 ± 0.1589 | 2.019 ± 0.1475 |
| fAHP (mV) | -12.30 ± 1.807 | -13.16 ± 1.994 |
| mAHP (mV) | -35.28 ± 1.994 | -38.25 ± 2.320 |
| Rin (mΩ) | 386.3 ± 62.24 | 495.6 ± 68.72 |

## Notes

### Competing Interest Statement

The authors have declared no competing interest.

